# A physiologically-based model of bile acid metabolism in mice

**DOI:** 10.1101/2022.11.10.515857

**Authors:** Bastian Kister, Alina Viehof, Ulrike Rolle-Kampczyk, Annika Schwentker, Nicole Simone Treichel, Susan Jennings, Theresa H. Wirtz, Lars M. Blank, Mathias W. Hornef, Martin von Bergen, Thomas Clavel, Lars Kuepfer

## Abstract

Bile acid (BA) metabolism is a complex system that includes a wide variety of primary and secondary, as well as conjugated and unconjugated BAs that undergo continuous enterohepatic circulation (EHC). Alterations in both composition and dynamics of BAs have been associated with various diseases. However, a mechanistic understanding of the relationship between altered BA metabolism and related diseases is lacking. Computational modeling may support functional analyses of the physiological processes involved in the EHC of BAs along the gut-liver axis. In this study, we developed a physiologically-based model of murine BA metabolism describing synthesis, conjugation, microbial transformations, systemic distribution, excretion and EHC of BAs at the whole-body level. For model development, BA metabolism of specific pathogen-free (SPF) mice was characterized in vivo by measuring BA levels and composition in various organs, expression of transporters along the gut and cecal microbiota composition. We found significantly different BA levels between male and female mice that could only be explained by adjusted expression of the hepatic enzymes and transporters in the model. Of note, this finding was in agreement with experimental observations. The model for SPF mice could also describe equivalent experimental data in germ-free mice by specifically switching of microbial activity in the intestine. The here presented model can therefore facilitate and guide functional analyses of BA metabolism in mice, e.g., the effect of pathophysiological alterations on BA metabolism and translation of results from mouse studies to a clinically relevant context through cross-species extrapolation.

## Introduction

Bile acids (BAs) are involved in many physiological processes in the body including digestion of nutrients or hormone metabolism [1,2]. The bile acid (BA) pool is a complex mixture of different BA species. Primary BAs are synthesized from cholesterol in the liver and are converted to various secondary BAs by the intestinal microbiome [3]. Both primary and secondary BAs are furthermore conjugated with either glycine or taurine in hepatocytes.

Within the body, BAs continuously undergo enterohepatic circulation (EHC) between the liver and the intestine. Hepatic BAs are secreted into the bile canaliculi and accumulate in the gallbladder. Upon food intake, gallbladder contractions release large amounts of the stored BAs into the small intestine to facilitate lipid absorption. BAs are further transported along the gut. Starting already within the small intestine but especially within the colon, BAs are subjected to microbial transformations such as deconjugation and the production of secondary BAs, e.g. by dehydrogenation. Most BAs are actively taken up by enterocytes, predominantly in the ileum, and further excreted towards portal blood. The remaining BAs are then either taken up by passive diffusion or they are secreted with the feces. From the portal blood, BAs are efficiently reabsorbed into the liver. Through sinusoidal transport they may subsequently reach the vascular circulation and eventually other tissues.

Due to the systemic nature of BA metabolism, diseases of both the liver (e.g. liver cirrhosis, liver cancer or inflammatory bowel disease) and the intestine (e.g. ulcerative colitis or Crohn’s Disease) have been associated with alterations in BA composition and distribution [2,4–11]. Such changes, however, may be difficult to investigate due to the complexity of the physiological processes involved and invasive sampling techniques required.

In this work, we developed a physiologically-based pharmacokinetic (PBPK) model of murine BA metabolism at the whole-body level which may be used as a platform for mechanistic investigation of BA metabolism. This model is of particular interest since mice are the most commonly used animal model to investigate human metabolism [12,13]. Mice produce cholic acid (CA) as well as muricholic acids (MCAs) that are made from chenodeoxycholic acid (CDCA). MCAs are hydroxylated at the C-6 position, which alters their physicochemical as well as the signalling properties. MCAs are more hydrophilic and less cytotoxic than other BAs and function as FXR antagonists instead of activating FXR signalling like other BAs [14]. Among others, this complicates translation of insights generated in mouse experiments to what can be expected in humans.

Our model describes the physiology of murine BA metabolism in great detail and can be used to simulate tissue concentration profiles of the most abundant BAs in mice. To inform the model, mice were experimentally characterized concerning their BA composition in various organs as well as their BA transporter expression along the gut axis and their cecal microbiota. Our model was further validated with complementary data set generated from germ-free mice. The here presented model can be used to simulate BA levels in tissues that are experimentally inaccessible. Likewise, it can be used to analyse the effect of pathophysiological alterations on BA metabolism. The model may hence serve as a tool for hypothesis testing and as a bridge between discoveries within mouse studies and clinical applications in human patients.

## Results

### A Physiologically-Based Model of Bile Acid Metabolism

The physiologically-based murine model of BA metabolism includes synthesis, hepatic and microbial transformations, (re-)circulation and excretion of BAs (Figure 1). For model development we used a PBPK model [15] in which BA metabolites were considered as the circulating molecules. The basic PBPK model represents the physiology of mice at a large level of detail. It therefore includes a significant amount of prior physiological knowledge regarding organ volumes, tissue composition, organ surface areas or blood perfusion rates [16,17]. Of note, the extrapolation to new scenarios and conditions is well possible due to the mechanistic structure of the underlying PBPK model [16,18].

**Fig 1.**
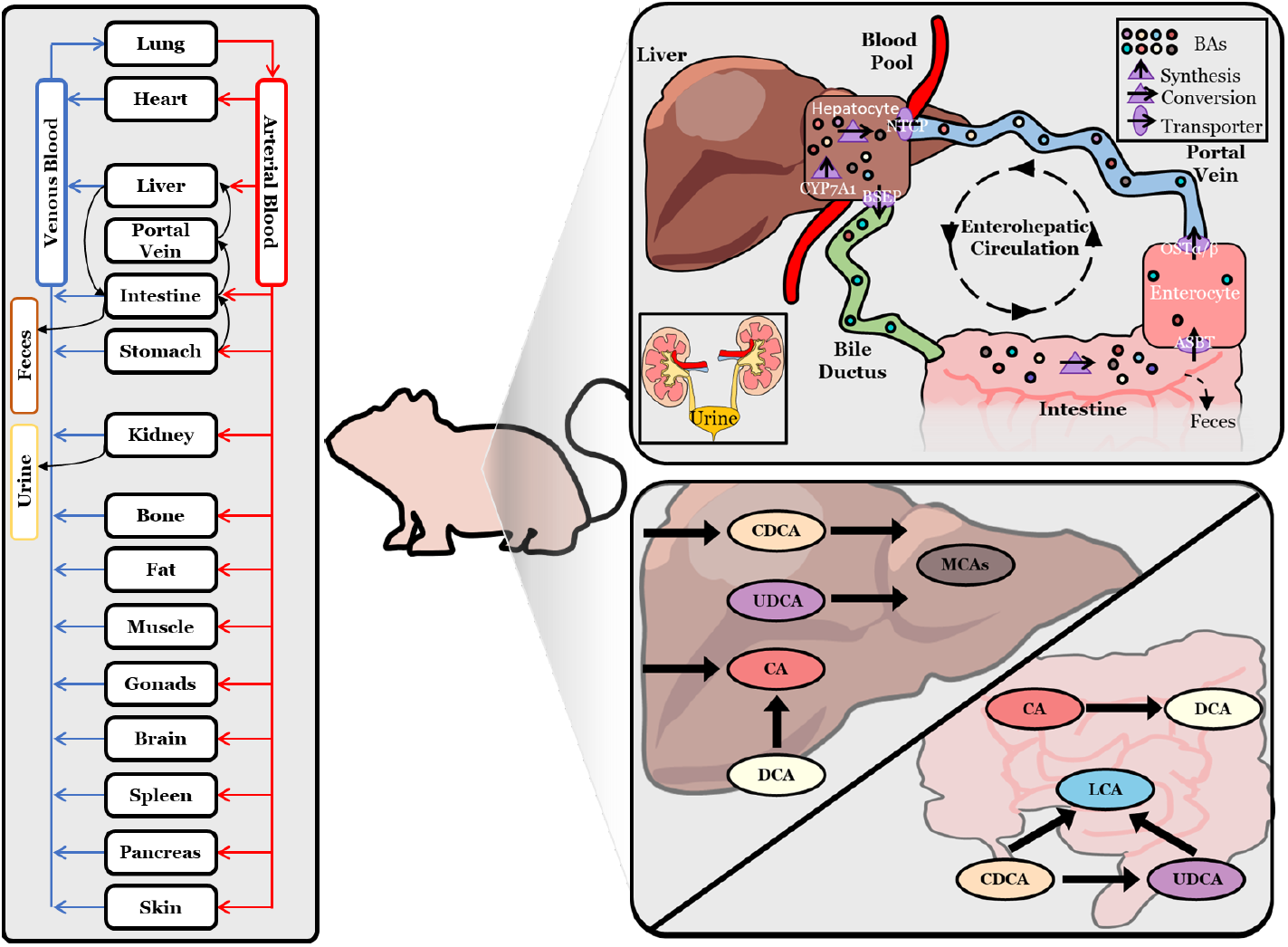
Physiologically-based bile acid model. Schematic overview of a PBPK model of bile acid biosynthesis via CYP7A1, hepatic and microbial transformation, active transport processes via BSEP, ASBT, OST-α/β and NTCP, as well as fecal and renal excretion. In the computational model, excretion to the gallbladder is neglected and BAs are directly secreted into the duodenum. Reactions of BAs are located either in the intracellular space of the liver or in the intestinal lumen.

In order to specifically inform physiological and kinetic parameters for bile acid metabolism in mice, extensive experimental data was collected. This data set comprised measurements of BA levels and composition in different tissues of specific pathogen-free (SPF) mice (Figure 2), physiological parameters (Figure 3), quantification of transporter gene expression in different segments of the gut as well as cecal microbiome diversity and composition (Figure 4).

**Fig 2.**
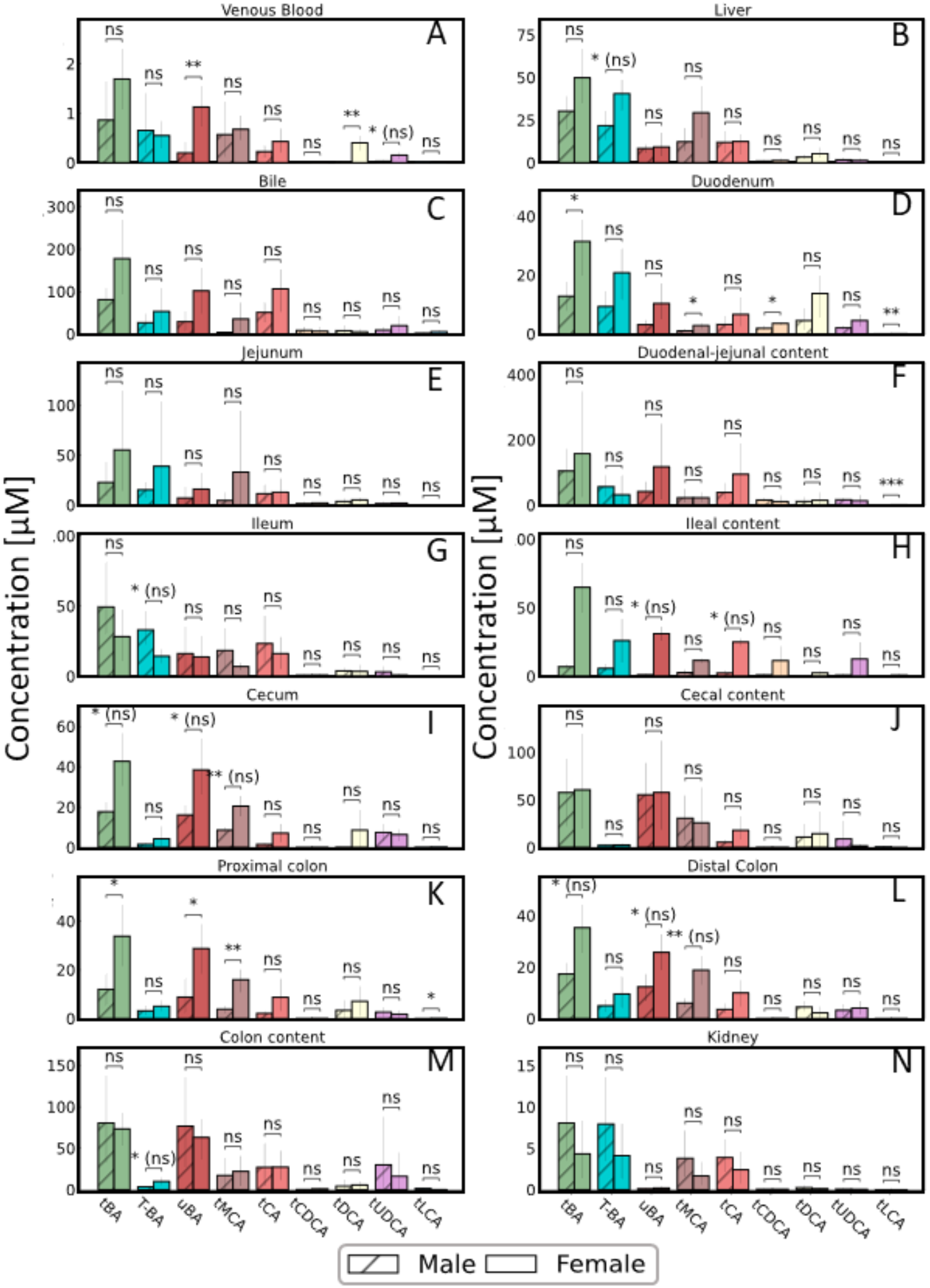
Bile acid levels in SPF mice. Concentration of total BAs (tBA), tauro-conjugated BAs (T-BA), unconjugated BA (uBA), total cholic acid (tCA), total muricholic acids (tMCA), total chenodeoxycholic acid (tCDCA), total deoxycholic acid (tDCA), total ursodeoxycholic acid (tUDCA) and total lithocholic acid (tLCA) in various organs and samples from male (striped bars) and female SPF mice (open bars). Statistical differences were assessed by independent t-test. Statistical significance is marked with asterisks and (ns) indicates non-significance after correction for multiple testing using Benjamini-Hochberg correction.

**Fig 3.**
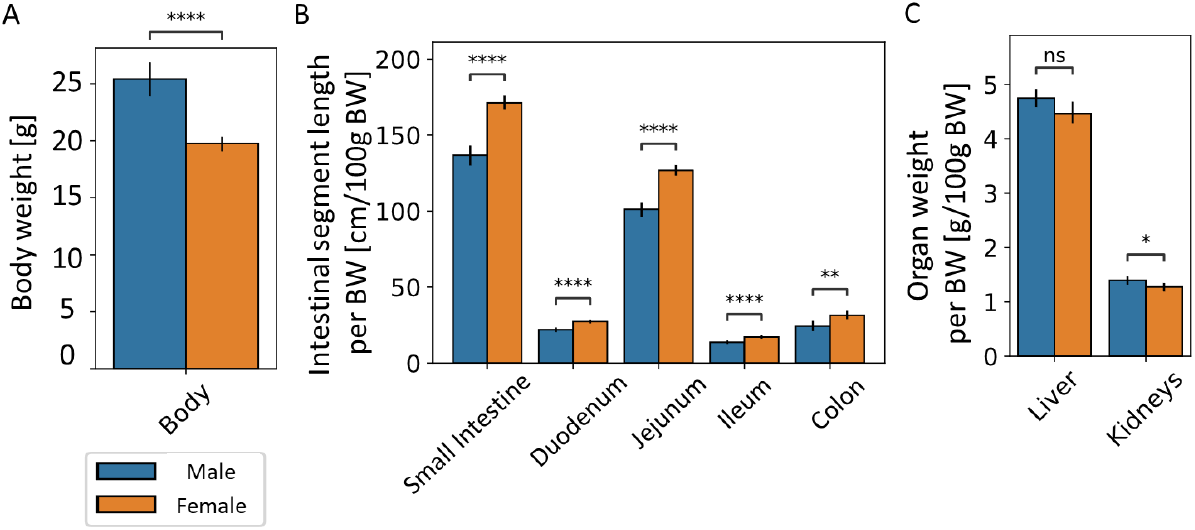
Physiological differences between male and female mice. Assessment of sex-related differences in body weight (A), length of intestinal segments (B) as well as weight of the liver and the kidneys (C) in SPF mice (male (blue bars), female (orange bars)). Significant differences were tested by two-way, independent t-test and significance was marked with asterisks.

**Fig 4.**
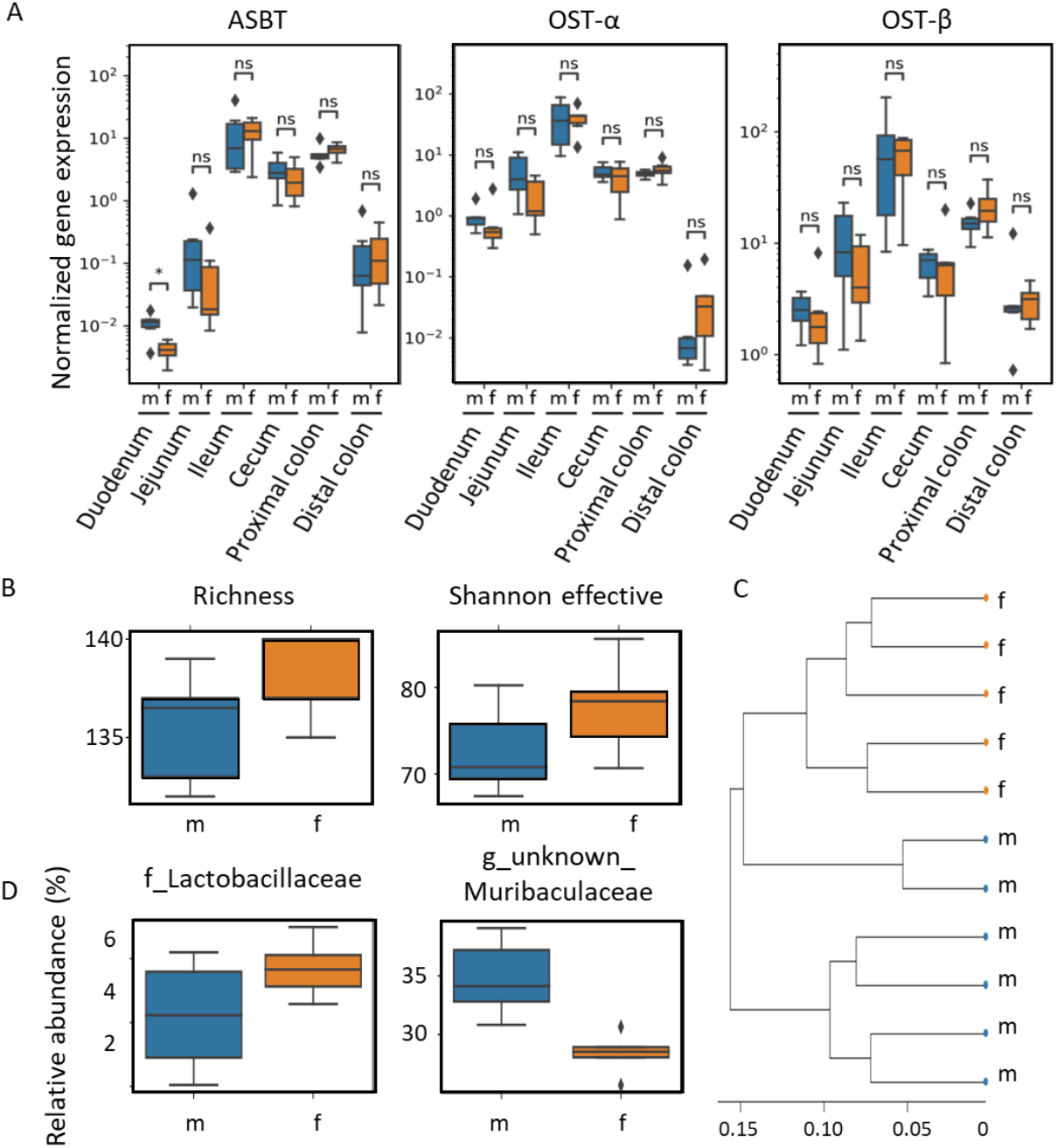
Sex-related differences in intestinal BA metabolism. Assessment of sex-related differences in the intestine relevant to BA metabolism: A) Expression of the BA transporters ASBT, OST-*α* and OST-*β* (from left to right) along the gut axis in male and female SPF mice measured by qPCR. Statistical significance was assessed by Mann-Whitney U-test and statistical significance is marked with asterisks (male (blue bars), female (orange bars)). B) Analysis of the cecal microbiome in male and female SPF mice by assessing observed species richness and effective Shannon index as indices of *α*-diversity, and C) *β*-diversity by hierarchical clustering of samples. D) The relative abundance of the family of Lactobacillaceae and an unknown genus of the Muribaculaceae, which significantly differed between male and female mice. Statistical significance was determined using Kruskal-Wallis and Wilcoxon rank sum test (α = 0.05).

An initial screening of BA levels showed that female mice had in general higher levels of BAs than their male littermates which is in agreement with published studies [19]. In our data, total BA concentration in female mice was consistently increased in venous blood serum, liver, bile and intestinal tissues (Figure 2A-E,I,K,L). Levels in the gut lumen were higher in the small intestine (Figure 2F,H), while the content of the large intestine did not show a clear picture (Figure 2J,M). Only in the ileal tissue, male mice displayed higher concentration of BAs than female mice (Figure 2G).

In order to account for sex-related variation in BA metabolism, we built separate physiologically-based models for male and female SPF mice. Using physiologically-based modeling to describe bile acid metabolism enabled us to incorporate explicit information on the organism’s physiology (Figure 3). In this study, male mice were approx. 30% heavier (Figure 3A), and correspondingly in female mice the liver and kidney were smaller (Figure 3C). Interestingly, the intestine had approximately the same length regardless of sex; thus, female mice showed a longer intestine in relation to their body weight (Figure 3B).

Four transport processes were considered in the model: (1) Excretion of BAs from the liver into the duodenum by the bile salt export pump (BSEP), (2) uptake from the gut lumen by the apical sodium-dependent bile acid transporter (ASBT), (3) excretion from enterocytes to portal blood by the organic solute and steroid transporter (OSTα/β) and (4) uptake of BAs from portal blood into hepatocytes by the sodium/taurocholate cotransporting polypeptide (NTCP). Expression of the transporters (ASBT, OSTα/β) along the gut axis showed overall no differences between male and female mice; however, they varied strongly between different gut segments (Figure 4A).

To examine potential differences in the microbiota of male and female mice, the microbial diversity and composition within the cecum was analyzed. First, we determined α-diversity for the within-sample taxonomic diversity. The determined species richness, as well as the Shannon effective counts, showed no significant sex-related difference (Figure 4B). Similarities in microbial community structure (β-diversity) were assessed based on generalized UniFrac distances [20,21]. We observed no significant separation of mice based on sex and an overall high similarity between all samples (Figure 4C).

In contrast, we found significant compositional differences in the cecal microbiome between male and female mice. In female mice, the family of *Lactobacillaceae* was more abundant, whereas an unknown genus within the family *Muribaculacea* was more prominent in male mice (Figure 4D). While various *Lactobacilla* species are able to metabolize BAs [22], the observed difference in relative abundance between male and female mice was only significant without p-value adjustment and overall rather low. For the other significantly abundant genus, no information is available linking it to BA metabolism. Overall, these results indicate that there are no major differences in intestinal transporter expression and microbiota diversity and composition between male and female mice.

Besides the aforementioned physiology of the organism, physicochemical properties such as molecular weight, solubility, lipophilicity (logP) and plasma-protein binding (fraction unbound) are a second important pillar of PBPK models [15]. Organ-plasma partitioning and passive transport can be directly derived from these parameters using an appropriate distribution model. For the physiologically-based model of bile acid metabolism, physicochemical properties of the tauro-conjugated forms (Table 1) were used to inform the compound properties of the PBPK model for small molecules.

**Table 1.** Physicochemical properties of bile acids used. Overview of physicochemical properties and their source that were used to inform compound specific parameters in the PBPK model. For the total BA (total CA: tCA, total CDCA: tCDCA, total UDCA: tUDCA, total DCA: tDCA, total LCA: tLCA, total MCAs: tMCA) the corresponding physicochemical values of the tauro-conjugated form (T-BA) were taken.

| BA species | Property | Value | Source |
| --- | --- | --- | --- |
| tCA | MW [g/mol] | 515.7 | PubChem Identifier: CID 6675 |
| tCA | Solubility [g/l] | 0.077 | ALOGPS (HMDB [28]) |
| tCA | logP | 0 | Heuman et al. [29] |
| tCA | pKa (acidic) | -0.88 | ChemAxon (HMDB [28]) |
| tCA | pKa (basic) | -0.053 | ChemAxon (HMDB [28]) |
| tCA | FU | 0.359 | Predicted [30] |
| tCDCA | MW [g/mol] | 499.7 | PubChem Identifier: CID |
| tCDCA | Solubility [g/l] | 0.00748 | ALOGPS (HMDB [28]) |
| tCDCA | logP | 0.46 | Heuman et al. [29] |
| tCDCA | pKa (acidic) | -0.99 | ChemAxon (HMDB [28]) |
| tCDCA | pKa (basic) | 0.18 | ChemAxon (HMDB [28]) |
| tCDCA | FU | 0.0776 | Predicted [30] |
| tUDCA | MW [g/mol] | 499.7 | PubChem Identifier: CID 9848818 |
| tUDCA | Solubility [g/l] | 0.0075 | ALOGPS (HMDB [28]) |
| tUDCA | logP | -0.94 | Heuman et al. [29] |
| tUDCA | pKa (acidic) | -0.99 | ChemAxon (HMDB [28]) |
| tUDCA | pKa (basic) | 0.18 | ChemAxon (HMDB [28]) |
| tUDCA | FU | 0.0776 | Predicted [30] |
| tDCA | MW [g/mol] | 499.7 | PubChem Identifier: CID 2733768 |
| tDCA | Solubility [g/l] | 0.0078 | ALOGPS (HMDB [28]) |
| tDCA | logP | 0.59 | Heuman et al. [29] |
| tDCA | pKa (acidic) | -0.75 | ChemAxon (HMDB [28]) |
| tDCA | pKa (basic) | -0.2 | ChemAxon (HMDB [28]) |
| tDCA | FU | 0.0768 | Predicted [30] |
| tLCA | MW [g/mol] | 483.7 | PubChem Identifier: CID 439763 |
| tLCA | Solubility [g/l] | 0.00028 | ALOGPS (HMDB [28]) |
| tLCA | logP | 1 | Heuman et al. [29] |
| tLCA | pKa (acidic) | -0.63 | ChemAxon (HMDB [28]) |
| tLCA | pKa (basic) | -1.1 | ChemAxon (HMDB [28]) |
| tLCA | FU | 0.0618 | Predicted [30] |
| tMCA | MW [g/mol] | 515.7 | PubChem Identifier: CID 168408 |
| tMCA | Solubility [g/l] | 0.075 | ALOGPS (HMDB [28]) |
| tMCA | logP | -0.81 | Heuman et al. [29] |
| tMCA | pKa (acidic) | -0.98 | ChemAxon (HMDB [28]) |
| tMCA | pKa (basic) | 0.084 | ChemAxon (HMDB [28]) |
| tMCA | FU | 0.365 | Predicted [30] |

In the computational model, total levels of CA, MCAs, CDCA, DCA, UDCA and LCA were considered as these represent the most abundant BAs that could also be measured in all compartments. De novo synthesis of BAs was considered as a constant formation rate in the intracellular space of the liver, and its magnitude was estimated from the excretion rate in feces [23–25] and urine [26]. Both excretion processes were considered by passive transport or active clearance, respectively. Subsequent formation of MCA using CDCA or UDCA as well as hepatic DCA hydroxylation was included in the model. Microbial metabolism of BAs was modeled as net enzymatic reactions, and the relative abundance of the corresponding enzymes along the gut was correlated with the activity of bile salt hydrolase (BSH) [27]. The included reactions were dehydroxylation of CA to DCA, and CDCA and UDCA to LCA as well as epimerization of CDCA to UDCA (Figure 1).

### Model calibration to data from SPF mice

For parameter estimation, we allowed sex-related differences in active hepatic processes. It was found that downregulation of BA synthesis and the transporter NTCP is both necessary and sufficient to explain BA composition and levels in male and female SPF mice (data not shown). This agrees with earlier findings showing upregulation of BA synthesis (Cyp7a1, Cyp27a1) [31] as well as an elevated expression of the basolateral uptake transporter NTCP in female mice [32]. This striking agreement with earlier findings is a first indication of the predictive capabilities of the computational model and generates confidence for further analyses.

Following parameter estimation, the final model adequately described BA levels in various organs, in both male and female SPF mice (Figure 5A-B). Up to 60% of the experimental data were recapitulated within one standard deviation (SD), 92% within a two-fold SD, and only 12 of 156 data points deviated by more than 2 SDs (Figure 5C-D). Even though the model describes a complex system and the measured data showed high variation, especially in the intestine, a good agreement between experimental data and model simulation was achieved.

**Fig 5.**
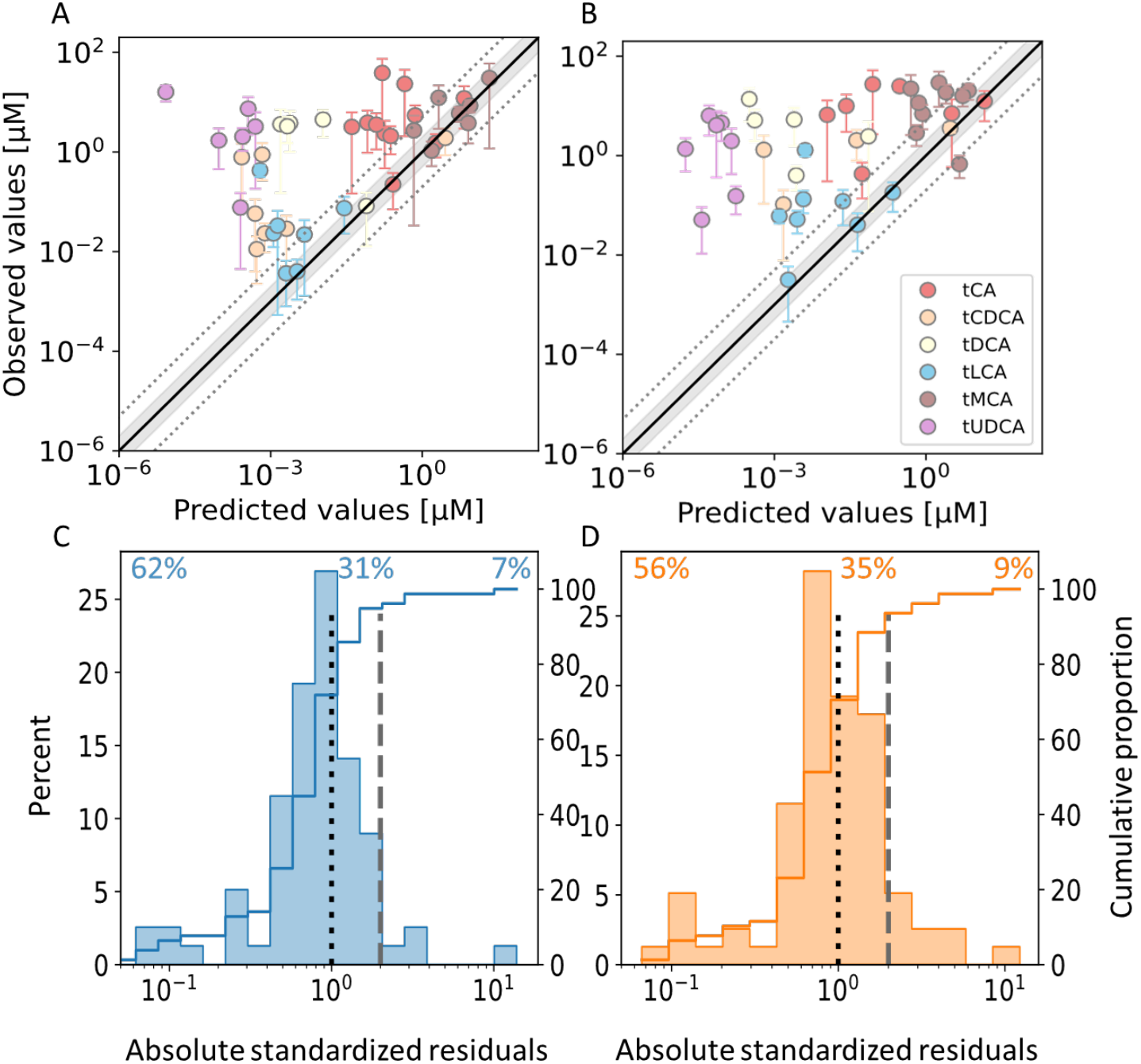
Model fit to data from SPF mice. Model simulations of bile acid concentration in male (A) and female mice (B) against corresponding data points used for fitting. Only data points with a coefficient of variation below 1 are shown. Unity is represented by a solid black line, a two- and five-fold range between predicted and observed values is indicated as a gray area or with dotted lines, respectively. Error bars show the SD. Distribution of the absolute standardized residuals between model simulations and data of male (C) and female (D) mice (histogram) and the corresponding cumulative function (line). The dotted and dashed lines indicate differences between model simulation and measured data of one SD and two SD, respectively. Cumulative proportions of predictions that are within one SD (top left, residuals left of dotted line), between one and two SD (top middle, residuals between dotted and dashed line) and above two SD (top right, residuals right of dashed line) of measured data are stated at the top of the panel.

### Model qualification to germ-free mice

Due to its mechanistic structure, the physiologically-based computational model of murine bile acid metabolism enabled the consideration of new scenarios. To validate the computational model of SPF mice, we next predicted BA levels in germ-free (GF) mice. In a first step, any microbial reaction in the SPF model was therefore removed, i.e. the corresponding rate was set to zero. Resulting predictions recapitulated BA concentration in male and female GF mice reasonably well (Figure 6A). About 59% of predicted concentrations fell within one standard deviation, 80% within a two-fold variation, and 15 predicted BA levels differed more strongly from the measured values (Figure 6B).

**Fig 6.**
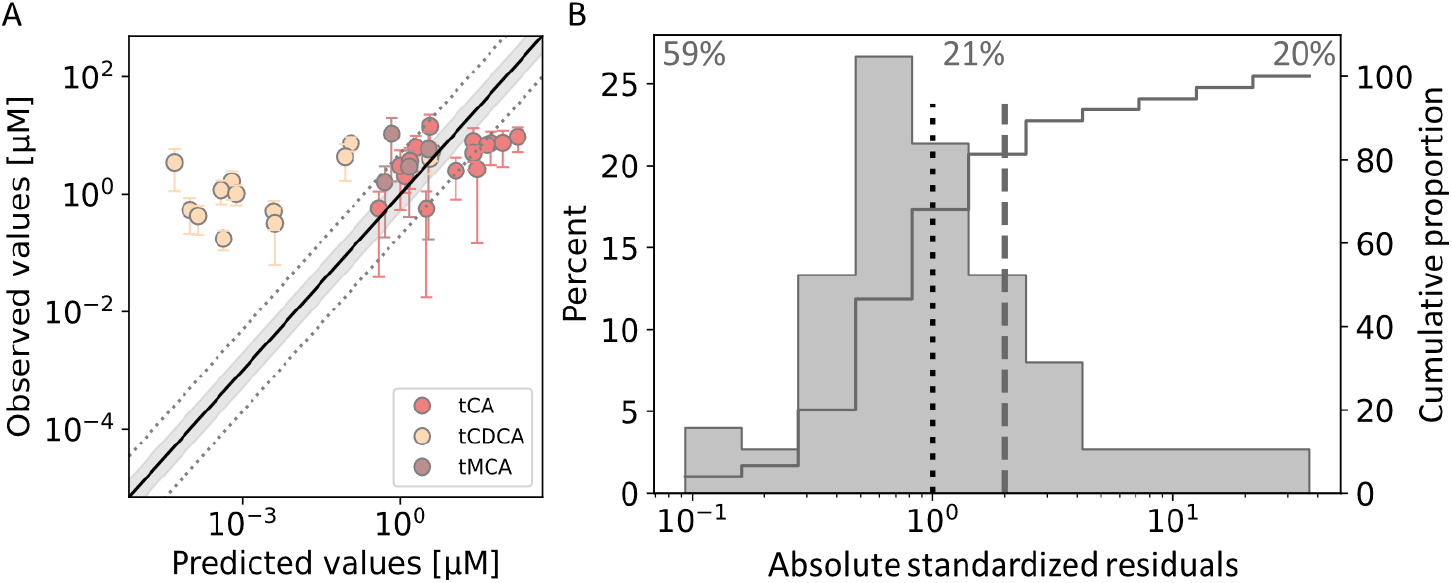
Model prediction of germ-free mice. Model predictions of concentration of bile acids in both male and female, germ-free mice against corresponding data points used for fitting (A). Only data points with a coefficient of variation below 1 are shown. Unity is shown as a solid black line shows unity, a two- and five-fold range between predicted and observed values is indicated as a gray area or with dotted lines, respectively. Error bars represent the standard deviation. B) Distribution of the absolute standardized residuals model predictions (histogram) and the corresponding cumulative function (line). The dotted and dashed lines indicate differences between model simulation and measured data of one SD and two SDs, respectively. Cumulative proportions of predictions that are within one SD (top left, residuals left of dotted line), between one and two SDs (top middle, residuals between dotted and dashed line) and above two SDs (top right, residuals right of dashed line) of measured data are stated at the top of the panel.

In a next step, we aimed to verify whether the inclusion of additional information about the physiology and intestinal transporter expression (Supplementary Figure S11 and S12) for GF mice would further improve the agreement between model simulations and experimentally measured BA concentrations. By doing so, model predictions worsened slightly. Less predictions could be recapitulated within one SD, but 80% were still within two SDs (Figure 7A) indicating that there are additional differences in BA metabolism between SPF and GF mice. Expression of the synthesizing enzymes CYP7A1, CYP27A1 and CYP7B1 were indeed shown to be elevated in GF mice [33–35]. However, increasing BA synthesis only yielded better model predictions by also allowing for differential regulation of other hepatic enzymes and processes. Upregulation of BSEP and downregulation of NTCP and MCA production from CDCA improved predictions only slightly (Figure 7B). The most accurate predictions of BA levels in GF mice could be obtained by allowing for reduced BA synthesis as well as differential regulation of hepatic processes. By doing so, 57% of predicted concentrations were within one SD, more than 80% within two SDs, and only 12% were not explained well.

**Fig 7.**
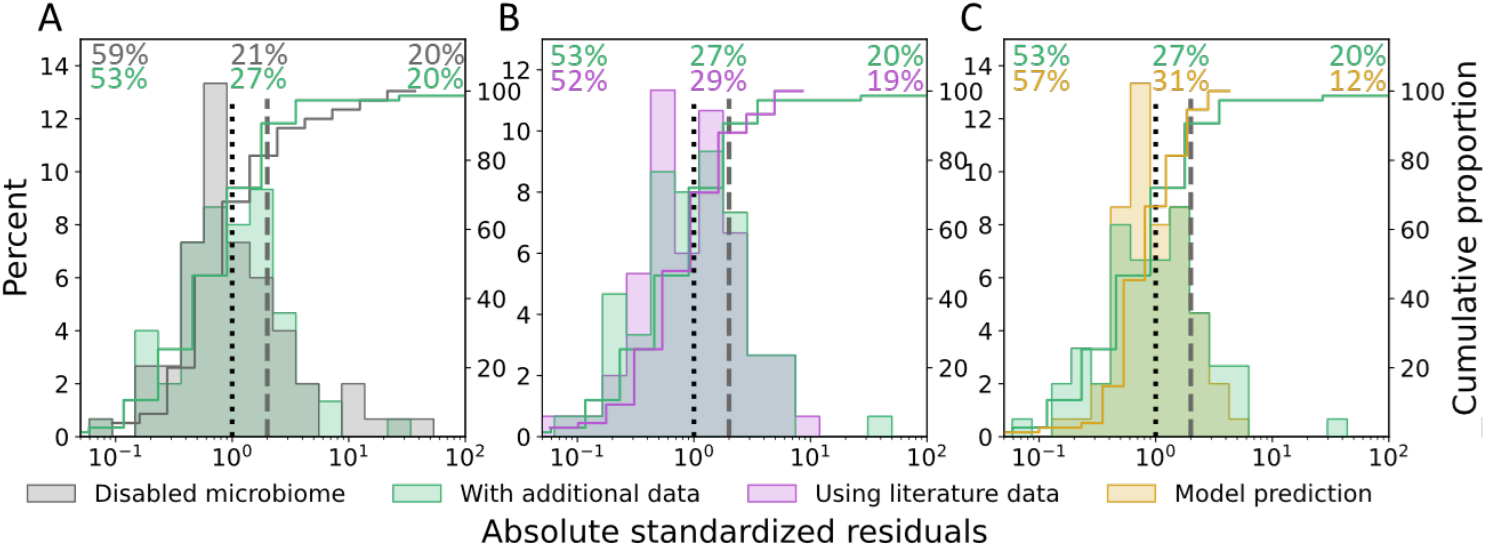
Model prediction of changes in germ-free mice. Comparison of different model variants for describing BA metabolism in germ-free mice by comparing the distribution of the absolute standardized residuals (histograms) and their corresponding cumulative function (lines). The dotted and dashed lines indicate differences between model simulation and measured data of one SD and two SDs, respectively. Cumulative proportions of predictions that lie within one SD (top left, residuals left of dotted line), between one and two SDs (top middle, residuals between dotted and dashed line) and above two SDs (top right, residuals right of dashed line) of measured data are stated at the top of the panel. A) Comparison of a simple extrapolation of the base model by disabling any microbial reaction (grey) against a model variant with additional information about physiology and intestinal transporter expression (green). The latter was also tested against model variants that introduce further expressional changes in the liver (B) according to literature (pink) or (C) as suggested by the model itself (yellow).

### Model analyses

After model calibration and validation, the here developed computational model described BA metabolism in both male and female mice with good agreement. As such, it could be applied to make comprehensive characterizations of the distribution and composition of BAs throughout the body (Figure 8). Model simulations showed highest BA levels within SI tissue, liver as well as intestinal lumen. Considerable BA amounts were also predicted within muscle, fat and skin. The levels are comparable to those found in organs of the EHC (Figure 8A, Supplementary Figure 13). The BA pool was estimated to undergo EHC 4.8 times per day in females and 4.1 times per day in males. Besides pool sizes, BA composition could be simulated even on sub-organ level. Composition was exemplarily assessed in liver, portal blood and skin in female mice (Figure 8B, Supplementary Figure 14). Model simulations showed that CA predominated in the liver, whereas in other organs MCAs constituted the most abundant BAs. The model was further used to simulate BA pools along the EHC and the gut axis (Figure 8C, Supplementary Figure 15). The model suggested a relative accumulation within the liver and the intestinal lumen, similar to other rodents where 70-95% of BAs are found in the lumen [36, 37]. Along the gut axis, bile acids accumulated strongly in the cecum and thus gave rise to higher levels throughout the LI. This coincides with the reduced transporter expression (Figure 4A) as well as intestinal transit rate (data not shown) in the cecum compared to the ileum.

**Fig 8.**
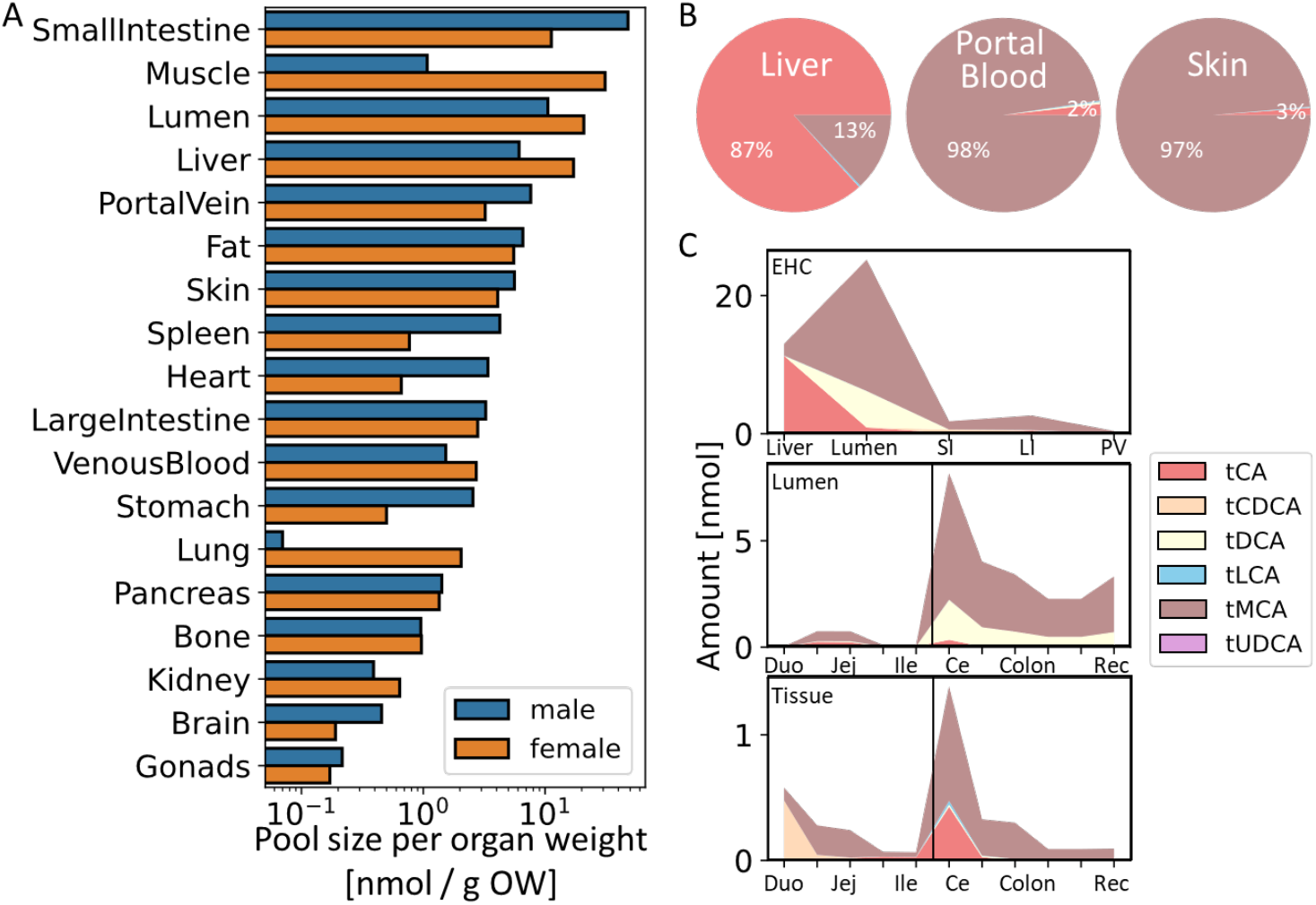
Model based BA pool distribution and composition at the whole body level. Simulated bile acid pool sizes per organ weight in different organs in male and female mice (A) and model-based bile acid composition in liver, portal blood plasma and skin in female mice (B) as well as simulated bile acid pools along the EHC and gut axis (C). For the EHC axis, BA level and composition are shown in liver, the intestinal lumen, small and large intestinal tissue (SI and LI) as well as portal blood plasma (PV). Along the gut axis, duodenum (Duo), jejunum (Jej), ileum (Ile), cecum (Ce), proximal and distal colon (Colon) and rectum (Rec) are shown and the transition of SI and LI are indicated by a vertical black line.

Apart from functional analyses of bile acid metabolism, the model could also be used to simulate physiological scenarios that had not been considered for model development. We here investigated in particular the functional effect of pathophysiological alterations on BA metabolism. To this end, the effect of bile acid malabsorption (BAM) as well as impaired intestinal barrier function on BA levels were simulated (Figure 9, Supplementary Figure 16). For BAM, impaired reabsorption in the terminal ileum (Figure 9A) as well as increased BA synthesis (Figure 9B) were considered [38]. Defective BA uptake by the ileal mucosa showed overall minor effects on BA pool sizes. Only within the LI, both tissue and lumen, increasing BA amounts could be observed. Increasing BA synthesis resulted in overall elevated BA levels, with the strongest effect on the LI and SI lumen and liver. Disruption of the intestinal barrier function depleted BA pools within the gut lumen, whereas in liver as well as venous and portal blood plasma BAs accumulated by 14- and 10-fold, respectively (Figure 9C). While the same trend was present in male mice, the accumulation of BAs was not as pronounced as in female mice showing a four- and three-fold difference, respectively (Supplementary Figure 16F).

**Fig 9.**
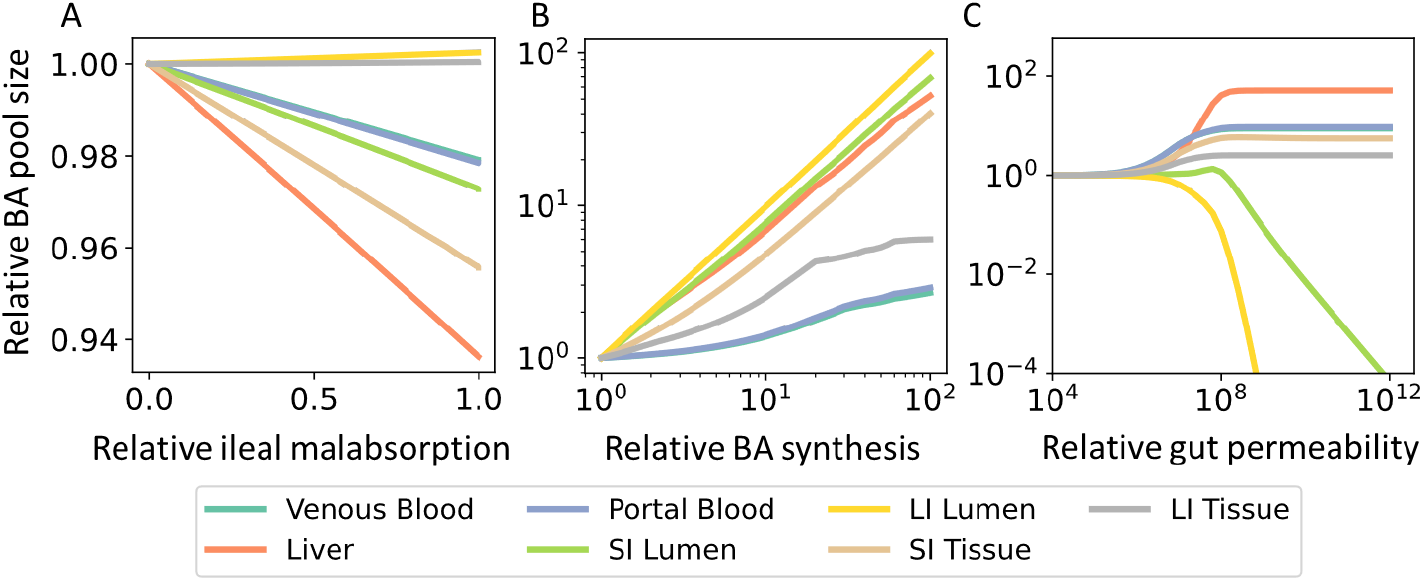
Model prediction of BA pool sizes in BA malabsorption and loss of intestinal barrier function. Predicted changes in bile acid pool sizes in different organs in female mice for decreasing BA absorption in the terminal ileum (A), for increasing BA synthesis (B) as well as increasing paracellular permeability along the whole gut (C).

## Discussion and Conclusion

In this work, we established a physiologically-based model of bile acid metabolism in mice. The model describes the systemic circulation, synthesis, hepatic and microbial conversions, and excretion of the most abundant BAs. It further addresses sex differences in BA concentration and composition that have been reported in the literature before but which were also prominent in our experimental data set. The model was carefully established and validated with an extensive data set specifically sampled from both male and female SPF mice. Thus, the model integrates and contextualizes heterogeneous data including BA concentration in different organs, transporter expression along the gut segments, physiological parameters and microbial composition in the cecum. Of particular note, the model suggested upregulation of BA synthesis and the transporter NTCP in female SPF mice. This observation, which is in agreement to earlier findings [31, 32], is an independent outcome of the model development process and had not been considered as prior knowledge for model building.

Furthermore, the resulting model for SPF mice could be used to predict BA measurements in GF mice by specifically eliminating microbial processes. This is a strong indication for the overall model quality and provides confidence that the model can be used for further analyses and predictions. The model was next extended with additional information specifying physiological parameters and intestinal transporter expression in GF mice. We found that upregulation of BA synthesis, as reported in literature [33–35], was not sufficient to explain BA levels in GF mice but had to be complemented by additional expressional changes in the liver. Unrestricted model predictions however suggested that BA synthesis might be downregulated. It remains to be investigated whether this discrepancy could be attributed to model inaccuracies or strain differences. This illustrates that the model can be applied for both extrapolation to unknown scenarios as well as contextualization of existing knowledge in a systemic manner, e.g., extrapolation to disease contexts or cross-species extrapolation.

Future analysis with the physiology based model may introduce structural revisions. In this version of the computational model, we included only the most abundant bile acid species but disregarded conjugation, sulfation and distinction of *α/β/ω*-MCA. Our model can therefore not capture the full complexity of the BA pool and might introduce a systemic bias in our predictions as different BA species do not have the same kinetics [39]. Furthermore, we had to simplify the dynamic behaviour of BA circulation. In the computational model, all BAs are secreted directly into the duodenum omitting the gallbladder. Thus, postprandial responses but also coprophagy, as a means of BA recycling [40, 41], were neglected as these effects are difficult to describe in mice. Consequently, the model cannot capture effects of the circadian rhythm as observed in other rodents [36, 37, 42, 43]. Moreover, the microbiota was only assessed in the cecum. The murine cecum is quite large and functions as a microbial fermentation vessel [44]; however, variation in the microbial composition in other gut segments might contribute to the observed sex differences. Previous studies have also shown that similar microbial communities might be enriched for different functions [45]. Besides compositional variation, there might be differences in microbial density between male and female mice which has not been assessed in this study. Despite these limitations, the model was able to recapitulate BA composition and levels at the whole body level and account for sex-related differences in BA metabolism, both with good accuracy.

Alterations in BA composition and dynamics have been associated with a plethora of diseases. Compositional changes have been reported for inflammatory bowel disease (IBD), ulcerative colitis, Crohn’s disease, liver cirrhosis, liver cancer, irritable bowel syndrome, short bowel syndrome, and obesity [2, 4, 5]. Impairments within the EHC of BA have been linked to cholestatic drug-induced liver injury, chronic liver disease, cholesterol gallstone disease, malabsorption, dyslipidemia and atherosclerosis [6–11]. In this context, the model was applied to predict the effect of BA malabsorption (BAM) as cause for idiopathic BA diarrhea (BAD) as proposed in [38] and impairment of the intestinal barrier function - as observed in celiac disease [46–49], IBD [50, 51] and metabolic associated fatty liver disease (MAFLD) [52–54] - on BA metabolism. Consistent with other studies, our model showed that an increase in BA synthesis [9, 55, 56] and overflow of reabsorption resulted in a strong accumulation of BAs in the LI lumen, whereas malabsorption in the terminal ileum without ileal or other obvious gastrointestinal disease did not suffice to induce BAD [57]. As a first assessment of a “leaky gut”, the paracellular intestinal permeability was altered in all gut segments equally. Further analyses may include other routes of transepithelial transport, i.e. transcellular or transporter-mediated, as well as regional differences in permeability as demonstrated in [58]. These predictions are possible since the model contains a detailed model of the gastrointestinal tract [59, 60] As demonstrated, the model can shed light on the complex interaction between pathophysiological alteration, such as physiological or expressional changes, microbial dysbiosis but also drug administration, and bile acid metabolism.

Investigating the link between BA metabolism and their role in human disease, various animal models have been applied, including lamprey, skate, zebrafish, rat, mouse, hamster, rabbit, prairie dog, and monkey [61–63]. Of the small animal models, hamsters are most similar to humans regarding BA metabolism [64–66]; nevertheless, mice remain the most commonly utilized animal model to investigate human metabolism [12,13]. Indicative of the difference between human and mice is the different bile acid composition [14,67]. The murine BA pool is heavily influenced by MCAs, increasing its hydrophilicity, lowering its cytotoxicity and shifting to a more antagonistic FXR-signalling regime [14]. Further differences relevant for BA metabolism can be found in the physiology of the GI tract [44,68–71], energy homeostasis [72] and the recycling of nutrients and bile acids through coprophagy [40, 41]. Therefore, extrapolation from mouse studies to humans for BA signalling or BA related diseases are difficult. The computational model developed in this work might support cross-species extrapolation due to the mechanistic structure of the underlying PBPK model [16]).

Lastly, the model can assist in optimizing experimental designs for mouse studies that aim to elucidate the complex behaviour of BAs in health and disease. This is especially relevant in the context of the “3R” principles proposed by Russel and Burch in 1959 [73]: Reduction, Refinement and Replacement of animal testing. In this context, we demonstrated that the model can predict BA level and composition throughout the body, most notably in experimentally not easily accessible organs, e.g. the liver or portal blood, under both physiological and pathophysiological conditions. Based on the same model predictions, the BA pool was estimated to be recycled four to five times per day in contrast to four to twelve times in humans [74]. This represents a first assessment of species differences between mouse and human. All of these predictions could be made without further animal sacrifice. We believe that the here presented model can serve as a useful platform for model-aided investigation of BA metabolism in prospective studies.

## Materials and Methods

### Mouse housing conditions and sampling

Samples were collected from germ-free (GF) and specific-pathogen free (SPF) C57BL/6N mice euthanized for scientific procedures in accordance with the German Animal Protection Law (TierSchG). The internal animal care and use committee (IACUC) at the University Hospital of RWTH Aachen approved the collection of gut content, body fluids, and organs from donor mice not subjected to any experimental treatment (internal approval no. 70018A4). GF mice were housed in isolators (NKPisotec, Flexible film isolator type 2D) under sterile conditions. To obtain SPF mice with C57BL/6N background, mice were taken from the isolator and colonised passively with a complex microbiota by cohousing with SPF mice. Mice of the first generations of C57BL/6N SPF mice after breeding were sampled for this work. Room temperature was kept between 21-24 °C and 25-40% humidity on a 12h:12h day:night cycle. All mice were fed a standard chow ad libitum (GF mice: *γ*-irradiated standard chow, V1124-927; SPF mice: autoclaved standard chow, ssniff V1124-300) and given autoclaved tap water (pH 7). Mice were housed in single sex cages with Tek-Fresh bedding (ENVIGO). Faecal samples of GF mice were taken to confirm the GF status via microscopic observation after Gram-staining and plating on both anaerobic and aerobic agar plates. Mice were sacrificed at an age of 12- to 13-weeks and blood, urine, gut tissue and content, liver, gall bladder and kidneys were collected. Systemic blood was collected from vena cava, put on ice for 5-10 min and subsequently centrifuged at 4,500 rpm for 15 minutes to obtain serum. The small intestine was divided by length into duodenum (proximal 16%), jejunum (middle 74%), and ileum (distal 10%). The colon was divided in proximal (50%) and distal (50%) parts. All samples got frozen immediately and stored at −80°C.

### Bile acid measurements

#### Sample preparation

First, x mg solid matrix were mixed with five times the *μ*l amount of ACN:water (1:1, v/v) and homogenised with a TissueLyser II (30 Hz, 10 min; Retsch Qiagen). After a short centrifugation (2 min, 14000 rpm) 100 *μ*l of the supernatant were added to 500 *μ*l ACN:water:methanol (3:1:2, v/v/v) and the sample was vortexed for 5 min. After sonication (5 min) and centrifugation (14,000 rpm, 4°C, 5 min), 550 *μ*l of the supernatant was transferred to a new tube and evaporated to dryness. The pellet was reconstituted in 100 *μ*l 50% and 10 *μ*L was used for analysis. For serum samples, 10 *μ*l samples were used.

#### LC-MS analysis

The analysis was performed using the validated Bile Acid Kit (Biocrates Life Sciences, Innsbruck, Austria) as described in Pham et al. [75]. For that 10 *μL* of the native samples/sample extract were pipetted onto a 96 well sandwich filter plate and prepared according to manufacturer’s instructions. For quantitation, 7 external calibration standards (each containing all 19 bile acids) and 10 isotope-labeled internal standards are used. A detailed list of metabolites is available at the manufacturer’s homepage Kit (Biocrates Life Sciences AG, Innsbruck, Austria). The LC-MS/MS analysis carried out by MRM acquisition using a Waters Acquity UPLC System coupled with QTRAP 5500 (AB Sciex, Concord, Canada). MP A consisted of 10 mM ammonium acetate and 0.015% formic acid, while MP B was of a mixture of acetonitrile /methanol/water (65/30/5;v:v:v), 10 mM ammonium acetate and 0.015% formic acid. Data processing is carried out with the provided quantitation method Kit (Biocrates Life Sciences AG, Innsbruck, Austria).

### Bile acid transporter expression

RNA isolation from homogenized tissue samples were performed using TRIzol reagent. Tissue homogenization was done using the FastPrep-24TM 5G from MP BiomedicalsTM. Isolated RNA was transcribed into cDNA using ReverseAid (Thermo Fisher) and RiboLock Inhibitor (Thermo Fisher). Quantitative PCR was done based on the use of taqman probes (Thermo Fisher) for the respective gene of interest (GOI). GOI expression was normalized to the expression of a housekeeping gene

### Microbiota analysis by high-throughput sequencing

#### Isolation of metagenomic DNA

For DNA isolation a modified protocol according to Godon et al. [76] was used. Frozen samples were mixed with 600 *μ*l stool DNA stabilizer (Stratec biomedical) and thawed at room temperature. After transfer to autoclaved 2-ml screw-cap tubes containing 500 mg of 0.1 mm-diameter silica/zirconia beads, 250 *μ*l 4 M guanidine thiocyanate in 0.1 M Tris (pH 7.5) and 500 *μ*l 5% N-lauroyl sarcosine in 0.1 M PBS (pH 8.0) were added. Samples were incubated at 70 °C and 700 rpm for 60 min. The cell disruption on a FastPrep® instrument (MP Biomedicals) fitted with a 24 × 2 ml cooling adaptor filled with dry ice was conducted 40 s at 6.5 M/s for 3 times. An amount of 15 mg Polyvinylpyrrolidone (PVPP) was added and samples were vortexed, followed by 3 min centrifugation at 15.000 x g and 4 °C. Approximately 650 *μ*l of the supernatant were transferred into a new 2 ml tube, which was centrifuged for 3 min at 15.000 x g and 4 °C. Of the supernatant 500 *μ*l were transferred into a new 2 ml tube and 50 μg of RNase were added before incubating for 20 minutes at 37 °C and 700 rpm. Subsequently gDNA was isolated using the NucleoSpin® gDNA Clean-up Kit from Macherey-Nagel according to the manufacturer’s protocol. DNA was eluted from columns twice using 40 *μ*l Elution buffer and concentration was measured with NanoDrop® (Thermo Scientific). Samples were stored at −20 °C.

#### Illumina sequencing of 16S rRNA gene amplicons

Library preparation and sequencing were performed as described in detail previously [77] using an automation platform (Biomek400, Beckman Coulter). Briefly, the V3-V4 region of 16S rRNA genes was amplified in duplicates (25 cycles) following a two-step protocol [78] using primers 341F-785R [79]. The AMPure XP system (Beckman Coulter) was used for purification before sequencing was carried out with pooled samples in paired-end modus (PE300) on a MiSeq system (Illumina, Inc.) with 25% (v/v) PhiX standard library according to the manufacturer’s instructions.

### Computational methods

#### PBPK modelling

Physiologically-based pharmacokinetic (PBPK) models describe the physiology of an organism at a large level of detail. Organs are explicitly represented in PBPK models and they are linked through systemic vascular circulation. Tissue concentrations can be simulated in PBPK models, even if they are experimentally inaccessible. Parameters in PBPK models explicitly represent specific physiological functions and they are taken from previously curated collections of parameters including organ volumes, surface areas, tissue composition and blood perfusion rates, respectively. For that reason, identification of PBPK models is limited to very few parameters, usually related to active processes underlying compound distribution of as well as elimination. PBPK models are hence based on a large amount of prior knowledge including detailed description of physiological processes such as enterohepatic circulation or absorption in the intestine

#### Software and calculations

The PBPK model of bile acid metabolism was established in PK-Sim® and further reactions and adjustments were done in MoBi® (Open Systems Pharmacology suite Version 11.150). Model simulations were performed using the ospsuite-R package in R (version 11.0.123). Parameter fitting was performed with the Monte Carlo algorithm implemented in the Open Systems Pharmacology suite. Residual calculation was set to linear and weights were derived from measured SD. Most reactions were defined as simple Michaelis-Menten kinetics, for BA synthesis a constant flux was assumed. Plotting and statistical testing was done with custom Python scripts. Where applicable, p-value correction for multiple testing was done using Benjamini-Hochberg correction using the statannotations package [80]. For assessment of impaired gut barrier function, unperturbed paracellular permeability of BAs was set to their corresponding transcellular permeability calculated by MoBi on basis of their physicochemical properties.

#### 16S rRNA amplicon data analysis

Data was analyzed with an updated version of a workflow previously described by Lagkouvardos et al. [77]. Raw reads were processed using the Integrated Microbial Next Generation Sequencing platform (www.imngs.org) [81] based on UPARSE [82]. For this, sequences were demultiplexed and trimmed to the first base with a quality score of at least 10. Subsequent pairing, chimera filtering as well as OTU clustering (97% identity) was performed using USEARCH 11.0 [83]. Sequences that had less than 350 and more than 500 nucleotides and paired reads with an expected error above 2 were excluded from further analysis. To avoid GC bias and non-random base composition, remaining reads were trimmed by fifteen nucleotides on each end. Clustering of operational taxonomic units (OTUs) was done at 97% sequence similarity. For further analysis, only those with a relative abundance above 0.25% in at least one sample were kept. Sequence alignment and taxonomic classification was done with SINA 1.6.1, applying the taxonomy of SILVA release 138 [84].

For assessment of microbial richness, diversity and community structure, the Rhea pipeline was used [85]. A detailed description of statistical tests applied are provided in the Rhea support information and in the corresponding scripts (https://lagkouvardos.github.io/Rhea). Normalization of sequence counts was done via simple division to their sample size and then multiplication by the size of the smaller sample before subsequent calculation of alpha-diversity parameters. Beta-diversity analyses were based on the calculation of unweighted and generalized UniFrac distances [20, 21].

## Acknowledgments

We thank Vanessa Baier and Zita Soons for the helpful discussions. We are grateful to Ntana Kousetzi (Functional Microbiome Research Group, Institute of Medical Microbiology, RWTH University Hospital) for outstanding technical support with microbiota analysis by sequencing. This manuscript has been released as a Pre-Print at bioRxiv [86].

## Funding

Funding was provided from the German Research Foundation (DFG) – Project-ID 403224013 – “SFB 1382”. MvB is thankful for support by Novo Nordisk Foundation grant NNF21OC0066551. MvB and URK are grateful for funding of the UFZ for the ProMetheus platform for proteomics and metabolomics.

## Author Contributions

**Bastian Kister:** Methodology, Software, Validation, Formal analysis, Writing–original draft, Writing–review & editing, Visualization **Alina Viehof:** Methodology, Investigation, Writing–review & editing **Ulrike Rolle-Kampczyk:** Investigation, Writing–review & editing **Annika Schwentker:** Investigation **Nicole Simone Treichel:** Formal analysis, Investigation, Writing–review & editing **Susan Jennings:** Investigation **Theresa H. Wirtz:** Writing–review & editing **Lars M. Blank:** Writing–review & editing **Mathias W. Hornef:** Resources, Writing–review & editing **Martin von Bergen:** Resources, Writing–review & editing **Thomas Clavel:** Resources, Writing–review & editing **Lars Kuepfer:** Conceptualization, Resources, Writing–original draft, Writing–review & editing, Supervision

## Data Availability

The raw amplicon sequencing data generated in the context of this study have been submitted to the European Nucleotide Archive and are available under project accession number PRJEB58856.

## Supporting Information

**Fig 10.**
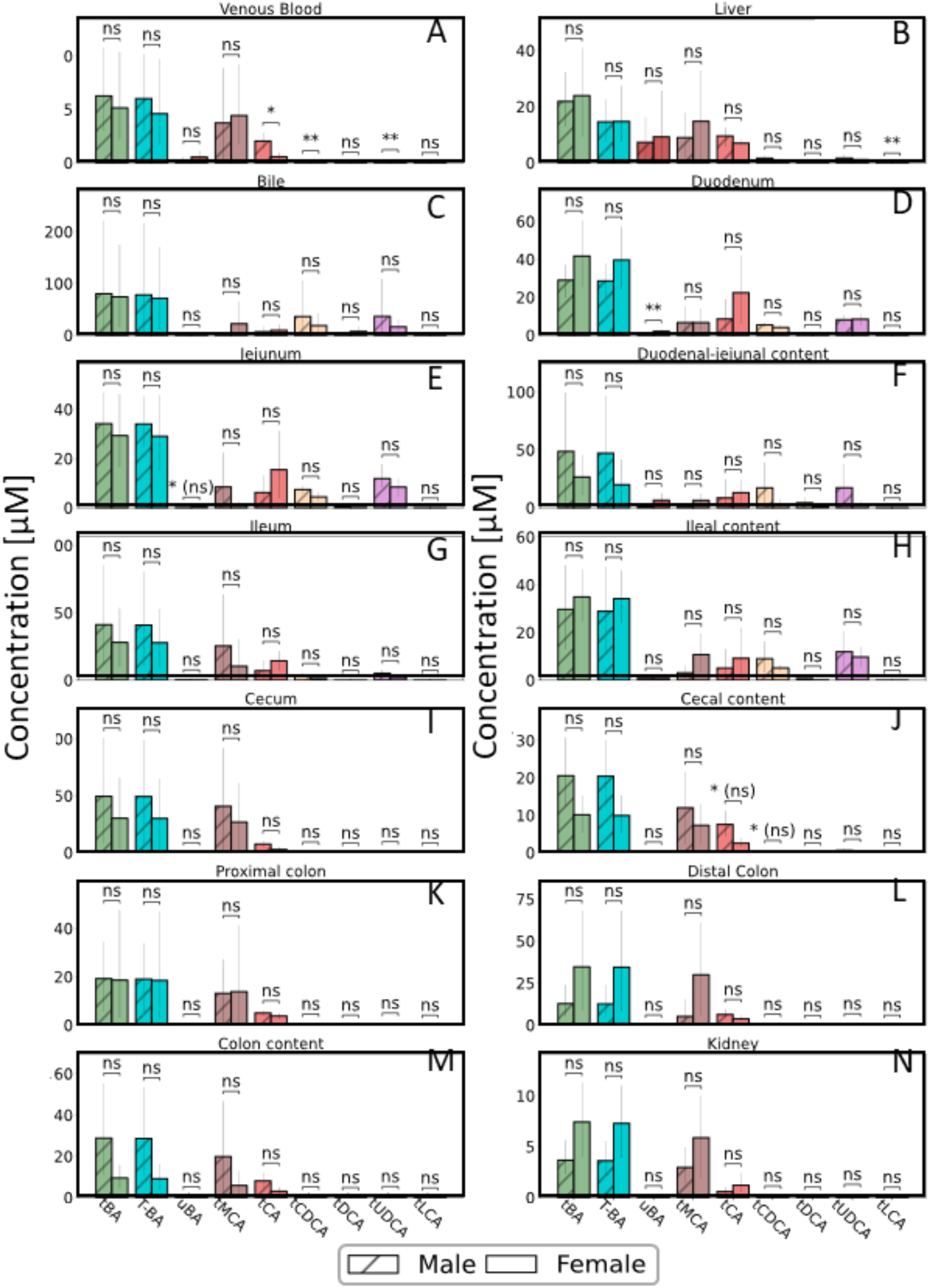
Bile acid levels in GF mice. Concentration of total BAs (tBA), tauro-conjugated BAs (T-BA), unconjugated BA (uBA), total cholic acid (tCA), total muricholic acids (tMCA) and total chenodeoxycholic acid (tCDCA) in various organs in male and female GF mice (saturated coloration). Statistical differences were assessed by independent t-test. Statistical significance is marked with asterisks.

**Fig 11.**
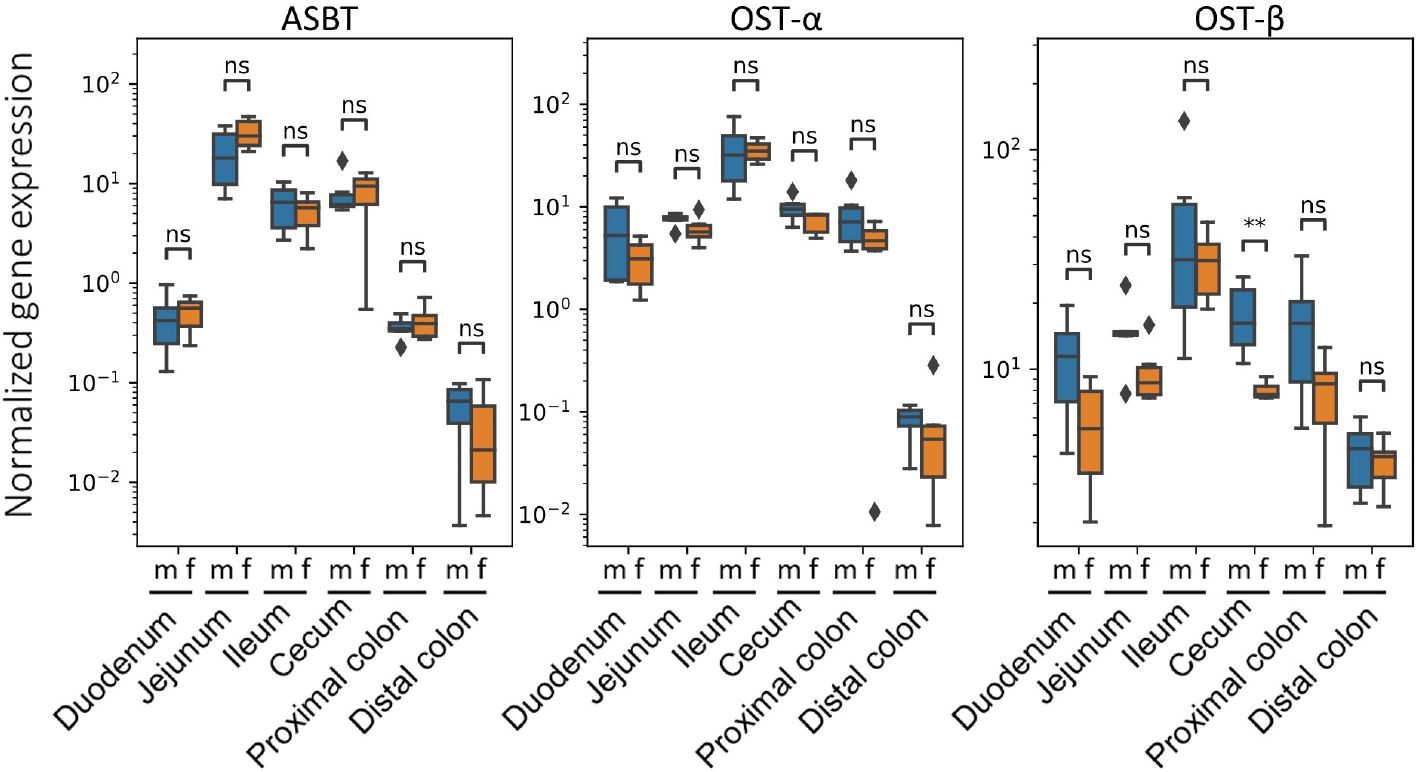
Sex-related differences in intestinal BA transporter expression. Assessment of sex-related differences in the expression of the BA transporters ASBT, OST-*α* and OST-*β* (from left to right) along the gut axis in male and female GF mice measured by qPCR. Statistical significance was assessed by Mann-Whitney U-test and statistical significance is marked with asterisks.

**Fig 12.**
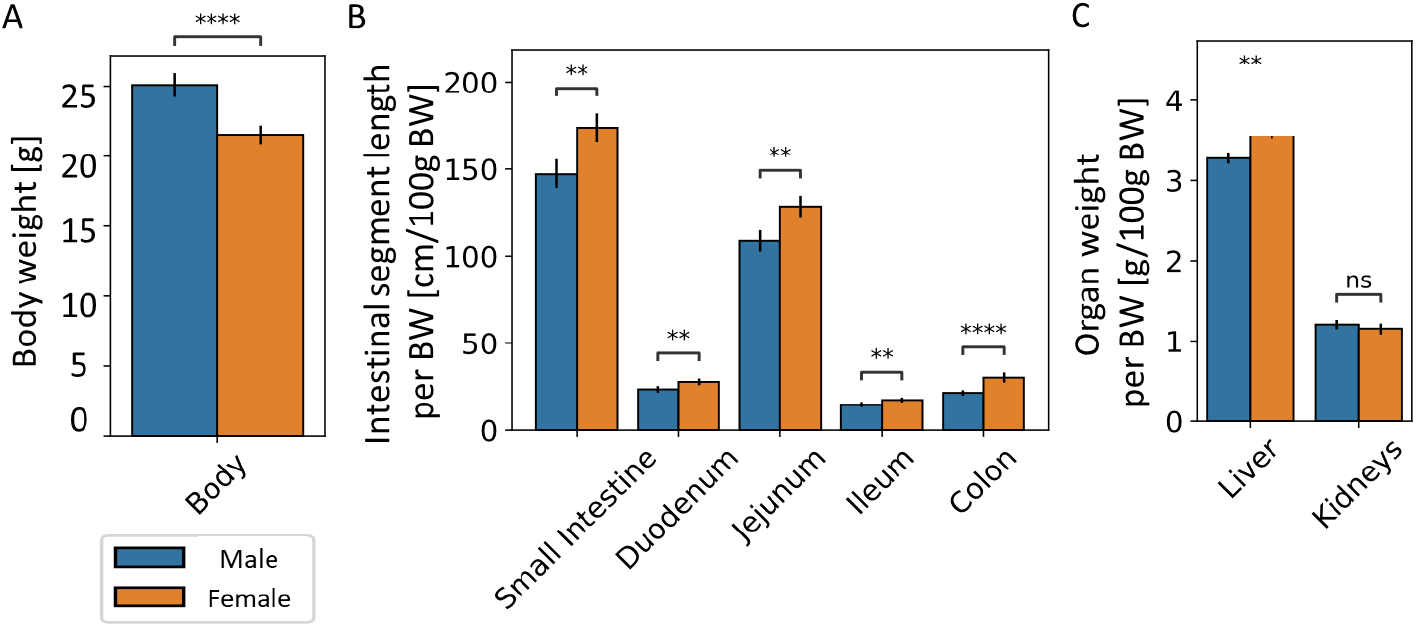
Physiological differences between male and female GF mice. Assessment of sex-related differences in body weight (A), length of intestinal segments (B) as well as weight of the liver and the kidneys (C) in GF mice. Significant differences were tested by two-way, independent t-test and significance was marked with asterisks and (ns) indicates non-significance after correction for multiple testing using Benjamini-Hochberg correction.

**Fig 13.**
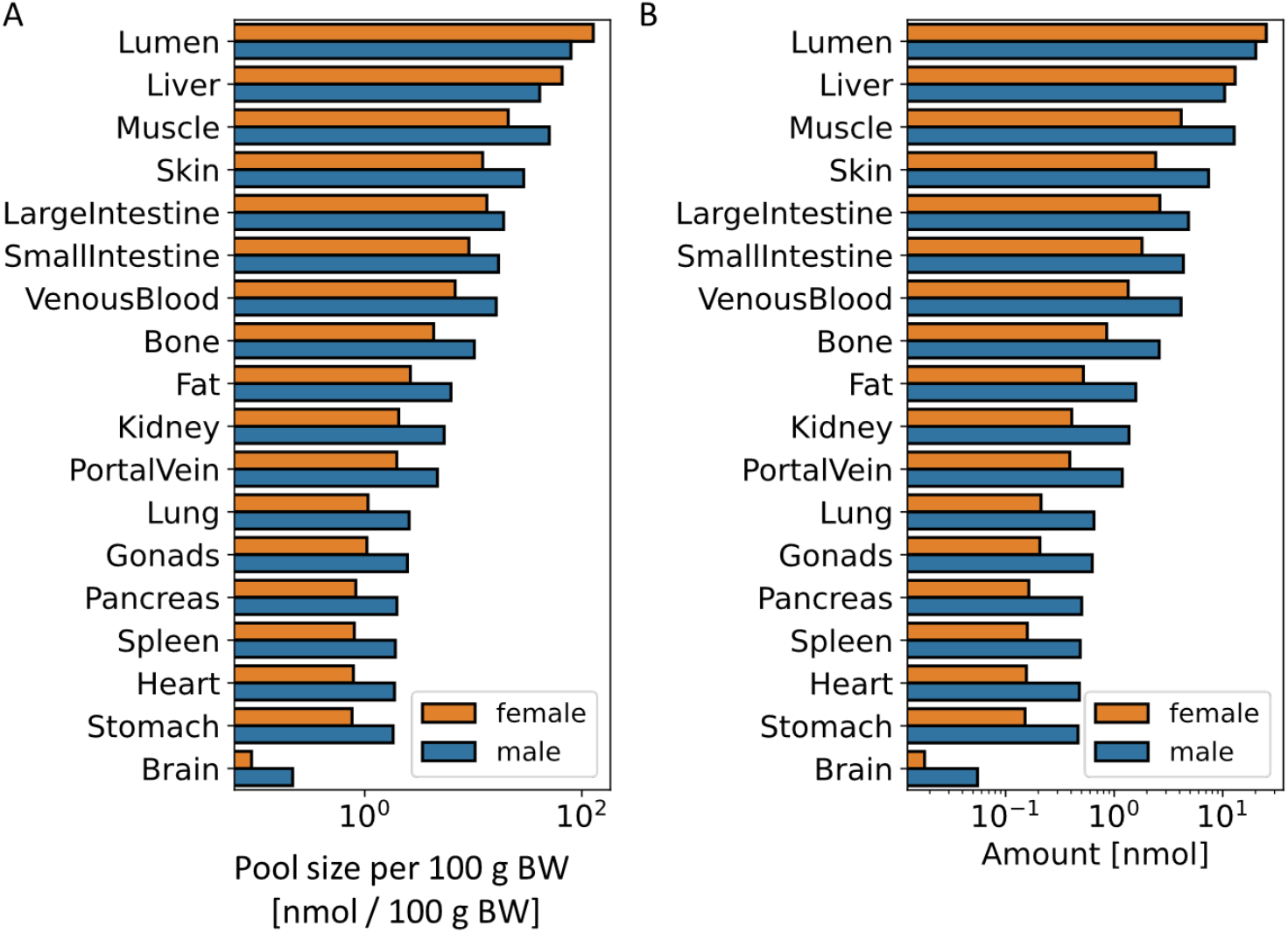
Model based BA pool distribution at the whole body level. Simulated bile acid pool sizes per body weight (A) and total BA pool sizes (B) in different organs in male and female mice.

**Fig 14.**
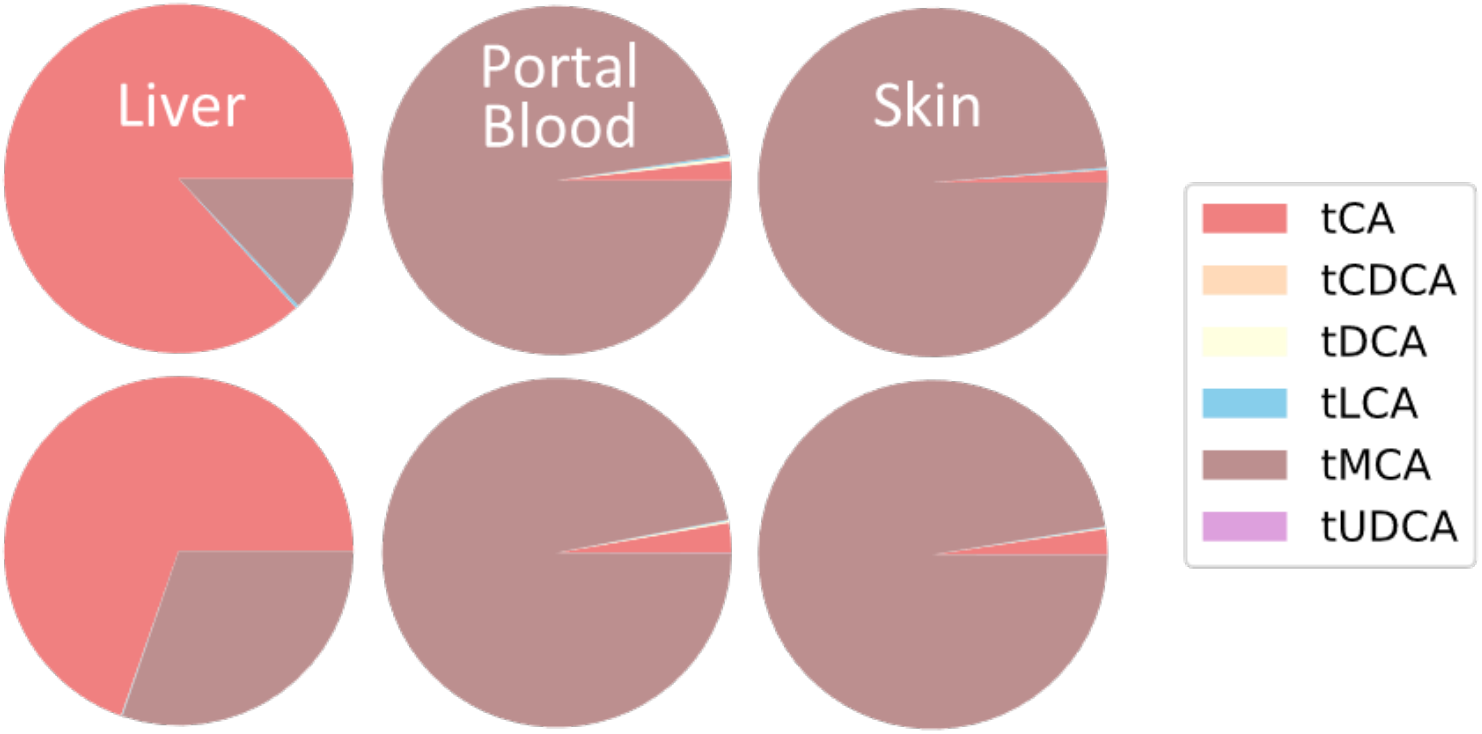
Model based BA pool composition in various organs. Simulated bile acid composition in liver, portal blood plasma and skin in female mice (top row)and male mice (bottom row).

**Fig 15.**
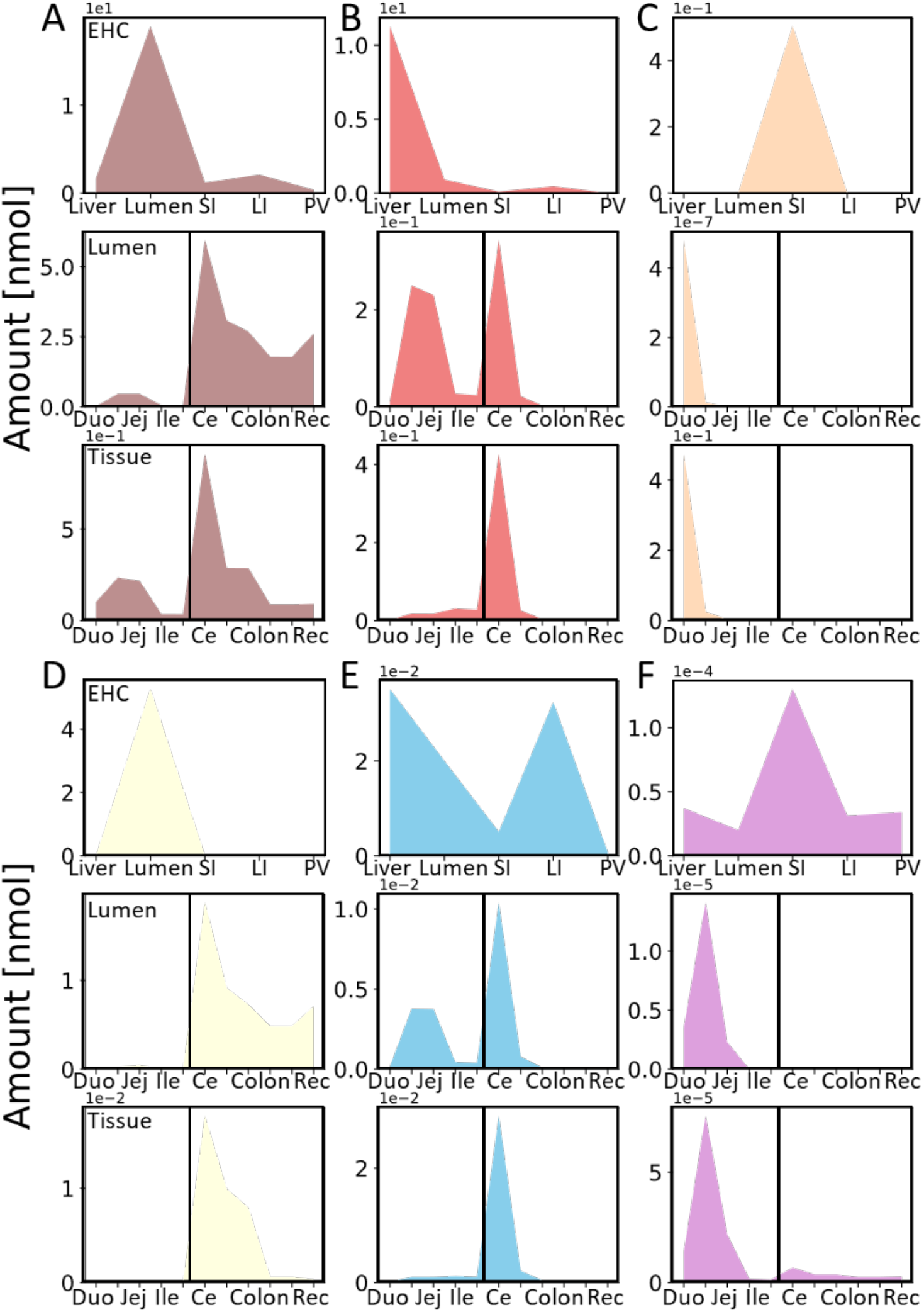
Model based BA pool distribution and composition along the EHC and gut axis. Simulated bile acid pools of total MCAs (A), total CA (B), total CDCA (C), total DCA (D), total LCA (E) and total UDCA (F) along the EHC and gut axis. For EHC axis, BA level and composition are shown in liver, the intestinal lumen, small and large intestinal tissue (SI and LI) as well as portal blood plasma (PV). Along the gut axis, duodenum (Duo), jejunum (Jej), ileum (Ile), cecum (Ce), proximal and distal colon (Colon) and rectum (Rec) are shown and the separation of SI and LI are indicated by a vertical black line.

**Fig 16.**
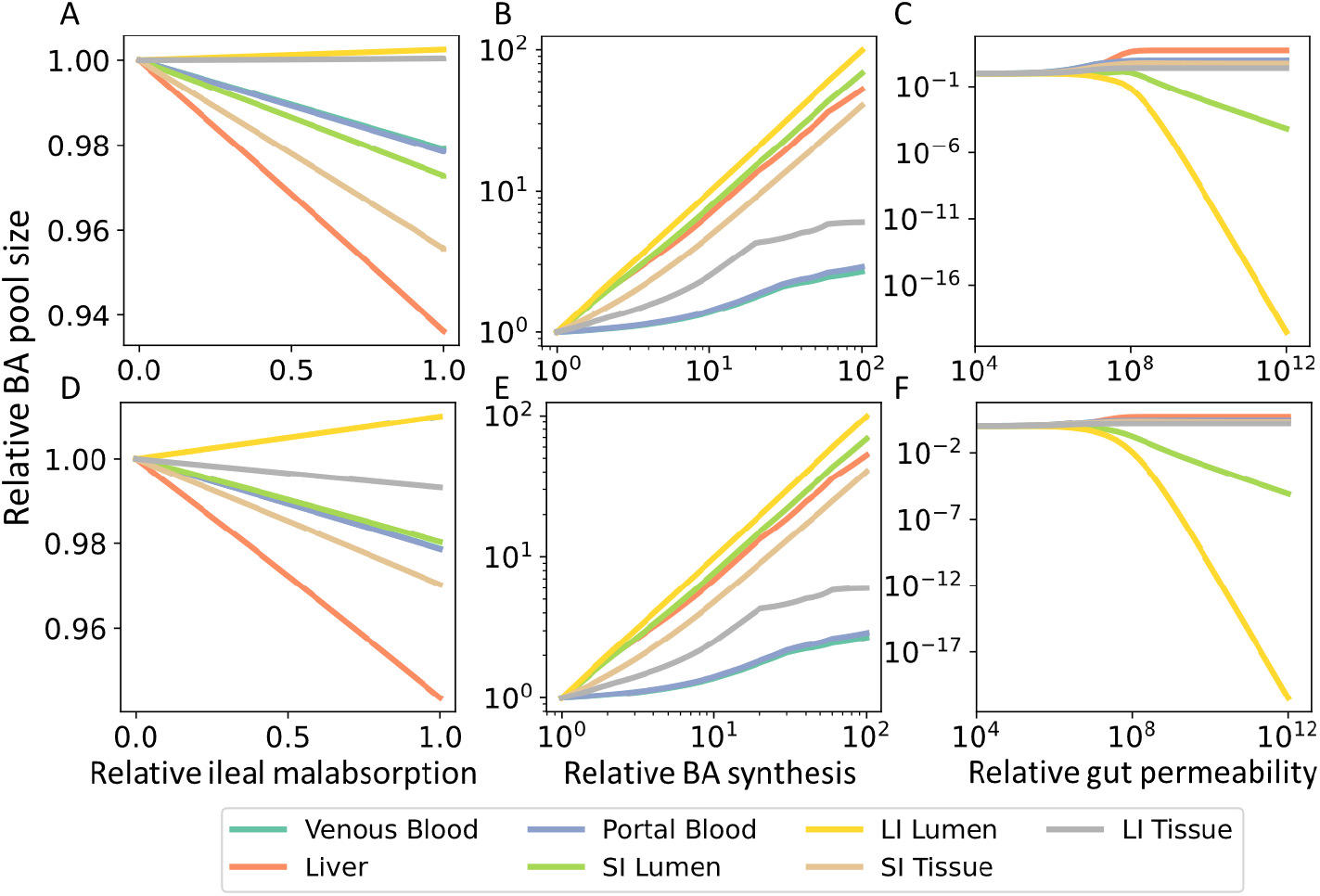
Model prediction of BA pool sizes in BA malabsorption and leaky gut. Predicted changes in bile acid pool sizes in different organs in female (top row) and male (bottom row) mice for decreasing BA absorption in the terminal ileum (A, D), for increasing BA synthesis (B, E) as well as increasing paracellular permeability along the whole gut (C, F).

## Notes

### Competing Interest Statement

The authors have declared no competing interest.

### Summary of Updates

A section on "Model analyses" including Figures 8 and 9 has been added,

